# Cortical circuit alterations precede disease onset in Huntington’s disease mice

**DOI:** 10.1101/391771

**Authors:** Johanna Neuner, Elena Katharina Schulz-Trieglaff, Sara Gutiérrez-Ángel, Fabian Hosp, Matthias Mann, Thomas Arzberger, Rüdiger Klein, Sabine Liebscher, Irina Dudanova

## Abstract

**Abstract:** Huntington’s disease (HD) is a devastating hereditary movement disorder, characterized by degeneration of neurons in the striatum and cortex. Studies in human patients and mouse HD models suggest that disturbances of neuronal function in the neocortex play an important role in the disease onset and progression. However, the precise nature and time course of cortical alterations in HD have remained elusive. Here, we use chronic *in vivo* two-photon calcium imaging to monitor the activity of single neurons in layer 2/3 of the primary motor cortex in awake, behaving R6/2 transgenic HD mice and wildtype littermates. R6/2 mice show age-dependent changes in neuronal activity with a clear increase in activity at the age of 8.5 weeks, preceding the onset of motor and neurological symptoms. Furthermore, quantitative proteomics demonstrate a pronounced downregulation of synaptic proteins in the cortex, and histological analyses in R6/2 mice and HD patient samples reveal reduced inputs from parvalbumin-positive interneurons onto layer 2/3 pyramidal cells. Thus, our study provides a time-resolved description as well as mechanistic details of cortical circuit dysfunction in HD.

*Significance statement:* Funtional alterations in the cortex are believed to play an important role in the pathogenesis of Huntington’s disease (HD). However, studies monitoring cortical activity in HD models *in vivo* at a single-cell resultion are still lacking. We have used chronic two-photon imaging to investigate changes in the activity of single neurons in the primary motor cortex of awake presymptomatic HD mice. We show that neuronal activity increases before the mice develop disease symptoms. Our histological analyses in mice and in human HD autopsy cases furthermore demonstrate a loss inhibitory synaptic terminals from parvalbimun-positive interneurons, revealing a potential mechanism of cortical circuit impairment in HD.

## Introduction

Huntington’s disease (HD) is an incurable hereditary neurodegenerative disorder, characterized by choreatic movements in combination with cognitive decline and psychiatric symptoms. HD is caused by an expansion of the CAG repeat in exon 1 of the Huntingtin gene (Huntington’s Disease Collborative Research Group, 1993), resulting in the expression of the aggregation-prone mutant Huntingtin (mHTT) protein with an elongated polyglutamine (polyQ) tract. mHTT interferes with multiple cellular functions, including transcription, energy metabolism, protein homeostasis and intracellular transport (Labbadia and Morimoto, 2013; Saudou and Humbert, 2016). The striatum is the most vulnerable region in HD, however, prominent pathological changes are also observed in the neocortex (Vonsattel and DiFiglia, 1998; Raymond et al., 2011; Waldvogel et al., 2012). Importantly, ample evidence points towards the disturbance of cortical function and impairment of corticostriatal communication as crucial early events in HD (Miller et al., 2011; Unschuld et al., 2012; Estrada-Sanchez and Rebec, 2013; Veldman and Yang, 2017). Imaging studies in HD gene expansion carriers demonstrate that cortical thinning and abnormalities of cortical activity contribute to the onset, progression and clinical variability of HD (Reading et al., 2004; Rosas et al., 2005; Rosas et al., 2008; Schippling et al., 2009; Nopoulos et al., 2010; Orth et al., 2010; Waldvogel et al., 2012). In particular, primary motor cortex (M1) is among the regions showing the earliest changes (Rosas et al., 2008), and the degree of cell loss in this area correlates with the motor impairments (Thu et al., 2010). In addition, analyses of tissue-specific conditional mouse models revealed the requirement of mHTT in both the striatum and the cortex for driving the full extent of HD phenotypes (Gu et al., 2005; Gu et al., 2007). Likewise, mHTT lowering in both regions is necessary for an efficient rescue of HD-related deficits (Wang et al., 2014; Estrada-Sanchez et al., 2015).

Cortical pyramidal neurons (principal cells, PCs) are known to be a vulnerable cell population in HD (Estrada-Sanchez and Rebec, 2013). HD mouse models display multiple morphological and electrophysiological abnormalities in these cells. Reduced dendritic arborizations and a decline in the density and stability of dendritic spines on PCs were observed in the somatosensory cortex (Klapstein et al., 2001; Murmu et al., 2013; Murmu et al., 2015). These defects were paralleled by lower levels of several synaptic proteins and a decrease in excitatory synapse density at an advanced disease stage (Murmu et al., 2015). In addition, electrophysiological recordings revealed changes in both excitatory and inhibitory inputs onto layer 2/3 (L2/3) PCs (Gu et al., 2005; Spampanato et al., 2008; Cummings et al., 2009).

Although the main focus in cortical HD pathology has been on PCs, there is also increasing evidence for an involvement of cortical interneurons (INs). Reductions in certain populations of INs were detected in human postmortem HD brains (Kim et al., 2014; Mehrabi et al., 2016). Furthermore, studies in conditional mouse models demonstrated the importance of mHTT expression in INs for the development of cortical pathology and behavioral defects (Gu et al., 2005), and attributed certain electrophysiological and behavioral alterations specifically to IN dysfunction (Dougherty et al., 2014). Despite these insights into the impairments occurring in PCs and INs, it has remained unclear how cortical network function is affected *in vivo* before disease onset and at different disease stages, and which molecular and circuit mechanisms underlie these functional alterations.

Here, we use chronic *in vivo* two-photon calcium imaging in awake, behaving HD mice to monitor the activity of large populations of L2/3 neurons in the M1 area at single-cell resolution. Our imaging experiments reveal disturbances of neuronal activity that precede the onset of symptoms. Proteomic analyses show a pronounced downregulation of synaptic proteins in the cortex, whereas histological findings in HD mouse brains and in human postmortem tissue point to a loss of parvalbumin (PV)-positive inhibitory synapses on PCs, providing a possible circuit mechanism for the cortical dysfunction in HD mice.

## Materials and Methods

### Transgenic mice

R6/2 mice (Mangiarini et al., 1996) transgenic for the 5′ end of the human *huntingtin* gene were maintained in a specific pathogen-free animal facility by crossing R6/2 males to F1 C57Bl6/CBA females. The presence of the transgene was verified by PCR with the following primers: forward, 5′-CCGCTCAGGTTCTGCTTTTA-3′, reverse, 5′-TGGAAGGACTTGAGGGACTC-3′. CAG repeat length was determined by Laragen and averaged 192 repeats. Mice were kept in inverted 14-10 h light-dark cycle from surgery on, had free access to food and water and showed no signs of inflammation or metabolic compromise. All procedures were performed in accordance with mouse protocols approved by the Government of Upper Bavaria (55.2-1-54-2532-168-2014, 55.2-1-54-2532-19-2015).

### Behavioral tests

#### Rotarod

Rotarod test was performed on a Rota-Rod NG (Ugo Basile). Mice were first trained on two consecutive days for 300 s at 5 rpm, and then tested on the accelerating rotarod from 5 to 40 rpm over a 300 s period. Latency to fall was recorded on three trials separated by 15 min resting periods, and the average value was taken for analysis.

#### Open field

Locomotor activity in the open field was assessed one hour after the beginning of the dark cycle. Mice were placed into a custom-made 40x40 cm arena, and total distance traveled was recorded for 10 min. The floor of the chamber was washed between the trials to minimize any olfactory cues that could affect exploratory behavior.

### Virus injection and cranial window surgery

For *in vivo* calcium imaging, an adeno-associated virus (AAV1/2) containing the genetically encoded calcium indicator GCaMP6s (Chen et al., 2013) and the bright structural marker mRuby2 (Lam et al., 2012) under the control of the human synapsin-1 promoter was used to label cortical neurons (Rose et al., 2016). Intracerebral injections of AAV and cranial window implantation were performed within the same surgery in 3.5-week-old mice deeply anesthetized with an intraperitoneal (i.p.) injection of ketamine/xylazine (130 and 10 mg kg^-1^ body weight, respectively). The analgesic carprofen (5 mg kg^-1^ body weight, subcutaneously) and the anti-inflammatory drug dexamethasone (10 mg kg^-1^, i.p.) were administered shortly before surgery. To increase viral uptake and spread, mannitol (20% solution; 30 ml kg^-1^ body weight, i.p.) was applied 20 min prior to virus injection (Burger et al., 2005). During surgery, the virus (titer: ~10^12^ infecting units per ml) was injected into L2/3 of M1 cortex (3 injection sites with stepwise 300 nl injections at 150, 200 and 250 μm depth). Next, a cranial window over the right cortical hemisphere was implanted as previously described (Holtmaat et al., 2009). Briefly, a circular piece of skull (4 mm in diameter) was removed over the fore- and hindlimb area of M1 (position: 1.3 mm lateral and 1.0 mm anterior to bregma) using a dental drill (Foredom). A round coverslip (VWR; d=4 mm) was glued to the skull using histoacryl glue (B.Braun) and dental acrylic cement (Kerr Vertise Flow). After surgery, mice received a subcutaneous injection of the antibiotic cefotaxime (60 mg kg^-1^) and were placed in a warm environment for recovery. After 10 days, a small custom-made metal bar (1 cm x 3 cm; 0.06 g) with a round opening was glued onto the coverslip with dental acrylic cement to allow for stable head fixation under the objective and repeated repositioning of mice during subsequent imaging sessions. Imaging began after a 21-day resting period after surgery.

### Handling and ball training

At the age of 5 weeks, mice were handled on 5 consecutive days for 10 min until they were familiarized with the trainer and routinely ran from hand to hand. In the subsequent ball training mice got adjusted to the experimental setup and head fixation. Ball training sessions that were repeated on 3 consecutive days were set in the dark (infrared (IR) light source) and lasted for 30 min. Mice were head-fixed by the metal bar to a custom-made holder and placed onto a styrofoam ball (d=20 cm) that floated on pressurized air and was custom-installed under the microscope objective (Dombeck et al., 2007). Mouse behavior (resting or running) was observed with an IR-sensitive camera (USB 2.0, 1/3″CMOS, 744x480 pixel; 8 mm M0814MP2 1.4-16C, 2/3′, megapixel c-mount objective; TIS) without exposure to additional stimuli or learning tasks. After the third session, mice had adjusted to the head fixation and showed alternating running and resting behavior.

### Two-photon calcium imaging in behaving mice

During the experiment, the same conditions as during ball training were applied. Mouse behavior was tracked at 15 Hz with an IR-sensitive video camera (TIS) and custom software (Input Controller, TIS). In addition, to track ball motion, a computer gaming mouse (G500S, Logitech) was positioned along the ball axis, controlled by a raspberry pi3 and custom written scripts using PuTTY to count ball rotation events (1000 counts/s). To synchronize *in vivo* two-photon imaging, behavioral video recording and ball speed measurements, a 900s lasting TTL pulse (5V) was sent to relevant hardware using Matlab (Mathworks). *In vivo* calcium imaging was performed with an upright multiphoton microscope (Bergamo II, Thorlabs) equipped with a Ti:Sapphire laser with dual beam (InSight DeepSee, Spectra Physics), a 8 kHz galvo/resonant scanner and a 16x, 0.8 NA water immersion objective (Nikon). The laser intensities were modulated with Pockels cells (Conoptics). The following wavelengths and emission filters were used to simultaneously image the two fluorophores: 920 nm / 525±25 nm (GCaMP6s) and 1040 nm / 607±35 nm (mRuby2). ThorImage 2.4 software (Thorlabs) was used for microscope control and image acquisition. To measure neuronal activity, time series images of selected positions/fields of view (FOVs) with bright expression of the calcium indicator were acquired at 10 Hz for a total duration of 900 s. For each mouse, two FOVs were acquired per imaging time point at the depth of 150-350 μm. Efforts were made to keep the GCaMP6s fluorescence constant throughout the entire experiment (<50 mW laser power out of objective). For each FOV, an image of the blood vessel map was acquired under epifluorescence light to ensure return to the same position in serial experiments. In addition, the information about XYZ-coordinates provided by the microscope stage was documented for each image position. The structural marker mRuby2 was used to precisely adjust the FOV in the z-plane. After imaging, the animal was returned back to its housing cage for rest. At the last imaging time point, the same FOVs were additionally imaged for 300 s during isoflurane (1.5%) anesthesia.

### Time series image processing and data analysis

Image analysis was performed with ImageJ (NIH) and Matlab software using custom written procedures. First, full frame images were registered and motion corrected in ImageJ using the moco plugin (Dubbs et al., 2016). Next, regions of interest (ROIs) were drawn manually around individual somata based on both maximum and mean intensity projections of all frames. For neighboring cells with direct contact, pixels containing signal from both neurons were excluded from the selection. For each imaging time point, the ROIs were visually inspected in the GCaMP and mRuby channel to ensure that the same cells were analyzed throughout the imaging period. The fluorescence intensity of all pixels inside each ROI was averaged and mean values were imported into Matlab for further processing as described previously (Liebscher et al., 2016). To account for neuropil contamination, the following correction method was applied: the initial ROI was fitted with an ellipse and this ellipse was stretched by 6 pixels. All pixels of the initial ROI, as well as those in neighboring ROIs were excluded from the resulting larger ellipse. Next, the corrected ROI signal was computed as follows: F_ROI_comp_ = F_ROI_ - 0.7 x (F_neuropil_ - median (F_neuropil_)) with F_ROI_comp_ representing the neuropil compensated fluorescence of the ROI, F_ROI_ referring to the fluorescence signal of the initial ROI selection and F_neuropil_ to the signal stemming from the neuropil (Liebscher et al., 2016). To estimate the baseline level (F0), each fluorescence trace was divided by the median of all values smaller than the 70^th^ percentile of the entire trace and subtracting 1 from those values, which reflects the baseline well as judged by visual inspection. Cells were classified as active in a particular experiment if they crossed a threshold of baseline + 3 x SD of the ΔF/F trace at least once for a minimum of 10 consecutive frames (1 s).

Video sequences acquired for behavioral tracking were analyzed in EthoVision (Noldus), using the activity analysis tool. Briefly, a ROI was drawn manually around the forepaws. Changes in pixels induced by forepaw movement were registered as activity change and plotted over time by determined algorithms. For direct comparison of video and imaging data at equal frame numbers, the 15 Hz videos were reduced to 10 Hz using the signal.resample function from Scipy and imported into Matlab for further analysis.

### MS data analysis and visualization

All data was obtained from the proteomic dataset published by Hosp et al. (2017). Principal component analysis (PCA) was conducted with the Perseus software package (Tyanova et al., 2016). Annotations were based on GOCC, GOBP, GOMF, CORUM, Pfam domains and KEGG pathway annotations with the exception of the main PCA drivers, which were complemented by manual annotations based on literature searches. All other analyses and data visualizations were performed with R software (R Development Core Team, 2008). Significantly changed protein expression was defined as a combination of a p-value lower than 0.05 and an expression change of at least two-fold compared to littermate controls.

### Immunofluorescent staining and confocal microscopy

Mice were transcardially perfused with phosphate-buffered saline (PBS) followed by 4% paraformaldehyde (PFA) in PBS. Brains were dissected out, post-fixed in 4% PFA at 4°C for 48 h, and coronal brain sections (70 μm) were cut on a microtome (VT 1000S, Leica). Free-floating sections were permeabilized in 0.5% TritonX-100 for 30 min and blocked in 5% normal donkey serum, 0.2% bovine serum albumin (BSA), 0.2% glycine, 0.2% lysine, 0.02% sodium azide in PBS for 2 h at room temperature, followed by overnight exposure to primary antibodies in 0.3% TritonX-100, 2% BSA, 0.02% sodium azide in PBS. The following primary antibodies were used: rabbit anti-PV, 1:500 (Abcam); mouse anti-NeuN, 1:500 (Millipore), guinea-pig anti-VGlut1 (Millipore), guinea-pig anti-vGlut2 (Millipore), mouse anti-PSD-95 (Sigma), mouse anti-Gephyrin (Synaptic systems), and rabbit anti-VGAT (Synaptic systems). After several washes with PBS, sections were incubated in corresponding Alexa secondary antibodies (Invitrogen) diluted 1:300 for 2 h followed by 10 min DAPI (Sigma) staining, several PBS washes and mounting with fluorescent mounting medium (DAKO). Fluorescence images were acquired with a Leica TCS SP8 confocal microscope using a 63x, 1.40 NA oil immersion objective (Leica). Image analysis was performed blindly with ImageJ and R. Briefly, images of pre- and postsynaptic stainings were converted into binary masks in ImageJ. Puncta of 2 pixels or less were excluded from further analysis. The coordinates of the remaining puncta were extracted using the “analyze particles” function. The distance of every presynaptic particle to every postsynaptic particle was then calculated using the Pythagorean equation and matrix calculations in R. A distance of 1 μm or less was counted as a synapse. For PV puncta quantification, the PC circumference was traced manually and measured with ImageJ. The area of the cell body and the surrounding PV staining were extracted and the PV-positive pixels counted using a custom written macro. The area of PV terminals around the cell body was normalized to the cell body perimeter.

### Patient material

5 μm paraffin sections from the primary motor cortex of 3 HD autopsy cases and 3 age-matched controls were obtained from Neurobiobank Munich, Ludwig-Maximilians University Munich. Demographic information is provided in Table 1. The experiments were approved by the local ethics committee. Sections were deparaffinized in Xylene twice for 5 min, hydrated in a decreasing ethanol concentration series, and transferred to warm tap water for 5 min. Antigen retrieval was conducted in boiling Tris-EDTA buffer (10 mM Tris, 1mM EDTA, 0.05 % Tween 20, pH 9.0) for 15 min. Slides were then transferred to tap water and blocked with BLOXALL Blocking Solution (Vector Laboratories) for 10 min and subsequently with 2.5 % horse serum for 30 min. The following primary antibodies were used: rabbit anti-PV, 1:250 (Abcam) and mouse anti-CamK2α, 1:250 (Abcam). After several washes with PBS, sections were incubated in corresponding Alexa secondary antibodies (Invitrogen) diluted 1:300, and Neurotrace, 1:500 (Life technologies) for 2 h followed by 10 min DAPI (Sigma) staining, several PBS washes and mounting with fluorescent mounting medium (DAKO). Fluorescence images were acquired with a Leica TCS SP8 confocal microscope using a 63x, 1.40 NA oil immersion objective (Leica). PCs in L2/3 were identified by their triangular shape, the presence of a nucleolus seen with Neurotrace, and/or through Camk2α-positive staining. Image analysis was performed with ImageJ and R. For PV puncta quantification, the circumference of the pyramidal neuron was traced manually, dilated by 1.5 μm and measured with ImageJ. The area of the cell body and the surrounding PV staining was extracted and the PV-positive pixels counted using a custom written macro. The area of PV terminals around the cell body was normalized to the cell body perimeter.

**Table 1.** Demographic data of HD patients and controls.

| Case no. | Sex | Age at death |
| --- | --- | --- |
| HD 1 | Male | 70 |
| HD 2 | Male | 60 |
| HD 3 | Male | 56 |
| Ctrl 1 | Male | 62 |
| Ctrl 2 | Male | 70 |
| Ctrl 3 | Female | 60 |

### Experimental design and statistical analysis

Female R6/2 mice and female wildtype (WT) littermates were used in all experiments. Behavioral tests were conducted with groups of 7-10 WT and 10 R6/2 mice. For chronic imaging experiments, 6 WT and 5 R6/2 mice were used. Proteomic data was from 4 WT and 4 R6/2 5-week-old mice and 3 WT and 3 R6/2 8-week-old mice. Immunostainings were performed on brain sections of 5 WT and 5 R6/2 mice at 5 weeks of age, and 5 WT and 4 R6/2 mice at 8 weeks of age. Human data was from 3 HD and 3 age-matched control autopsy cases. For behavioral tests, comparisons were made with two-way ANOVA with Bonferroni’s multiple comparison test. Two-way repeated measures ANOVA was used to reveal effects of genotype and time in chronic imaging experiments. Pearson’s Chi-square test was applied to evaluate changes in activity categories. Exact binomial two-sided test was used for binomially distributed data. Two-tailed unpaired Student’s t-test was applied for comparisons of two groups in histological analyses. Data were analyzed in a blinded manner. Data are expressed as mean ± SEM unless indicated otherwise, with p<0.05 defining differences as statistically significant (*p<0.05; **p<0.01; ***p<0.001; n.s. - not significant).

## Results

### Chronic two-photon calcium imaging in the cortex of R6/2 mice

We investigated neuronal activity in R6/2 transgenic mice, which express mHTT-exon 1 with a pathological polyQ expansion under the human HTT promoter and are characterized by an early onset and rapid progression of disease (Fig. 1A) (Mangiarini et al., 1996; Carter et al., 1999; Meade et al., 2002). In spite of considerable brain atrophy, no obvious cell death is observed in this line until the age of 12 weeks (Mangiarini et al., 1996; Dodds et al., 2014). Overt neurological symptoms such as tremor, dyskinesia and balance impairment start at 9-11 weeks of age (Mangiarini et al., 1996). The exact age of onset of motor impairments varies between different R6/2 colonies and is dependent on the CAG repeat length. Surprisingly, very high repeat numbers, which occur due to the genetic instability of the repeats, lead to a later onset and overall attenuation of the HD-related phenotypes (Morton et al., 2009; Cummings et al., 2012). The CAG repeat length in our colony averaged 192 ± 2 repeats and was higher than in the original line (~150 repeats) (Mangiarini et al., 1996). We therefore characterized the motor phenotype in our colony by testing the mice on an accelerating rotarod and in the open field. 8 to 9-week-old R6/2 mice showed no significant deficits on the rotarod and normal locomotion in the open field, whereas by the age of 12-13 weeks, they were severely impaired in both tests (Two-way ANOVA with Bonferroni’s multiple comparison test; Rotarod, 8-9 weeks, p>0.05; 12-13 weeks, p<0.001; Open field, 8-9 weeks, p>0.05; 12-13 weeks, p<0.001; Fig. 1B). We conclude that the onset of motor defects in our colony occurs after 9 weeks of age and is thus slightly delayed compared to other reports (Carter et al., 1999; Murmu et al., 2013), likely due to the expansion of CAG repeats and/or differences in housing conditions.

**Figure 1.**
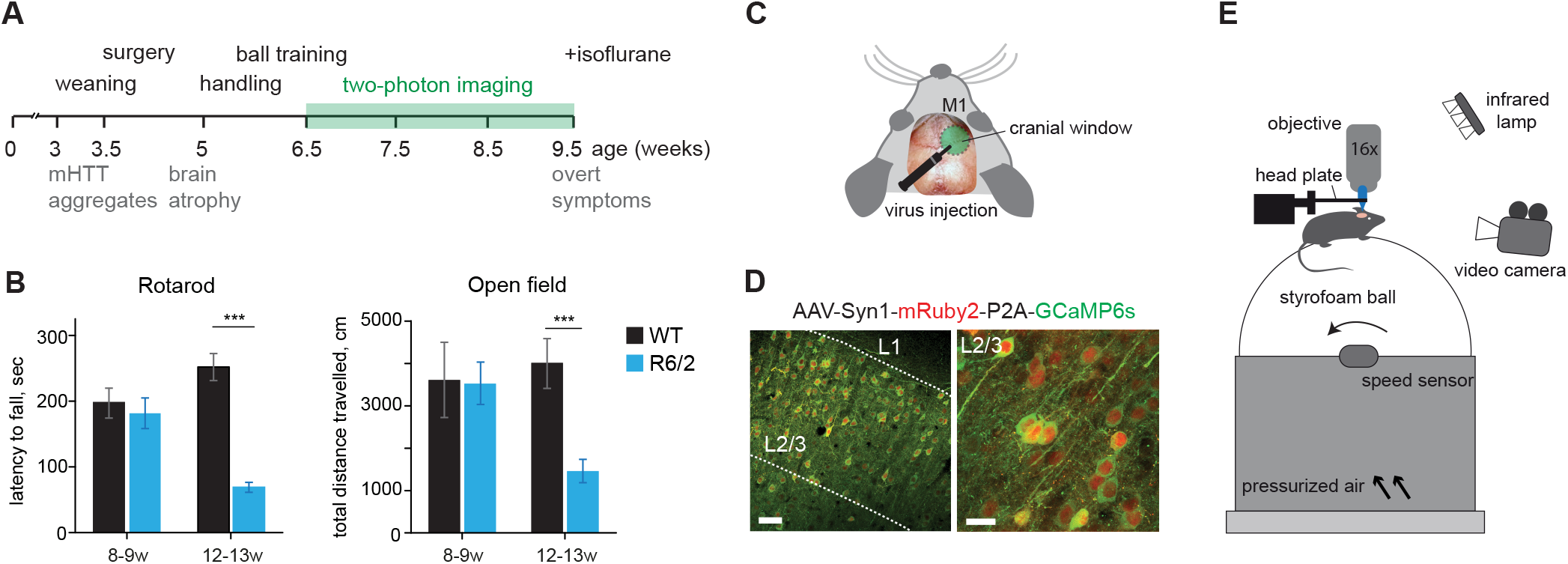
Chronic two-photon calcium imaging in R6/2 mice. **A**, Experimental design and timeline of R6/2 phenotypes. **B**, Left: Latency to fall on the accelerating rotarod. Right: Distance travelled in the open field. ***p<0.001. **C**, Scheme of a cranial window over the M1 cortex. **D**, Coronal brain section showing AAV-mediated (AAV1/2-Syn1-mRuby2-P2A-GCaMP6s) expression of mRuby2 (red) and GCaMP6s (green) in L2/3 neurons in M1. **E**, Imaging setup. A mouse head-fixed under a two-photon microscope is placed on a styrofoam ball floating on pressurized air. Neuronal activity is monitored through a cranial window. Running behavior is registered by an IR-sensitive video camera and a speed sensor. Scale bars: D, left image, 50 μm; right image, 20 μm.

To study longitudinal changes in neuronal function before and around the onset of disease, we chronically monitored calcium responses in L2/3 neurons in the M1 cortex of R6/2 mice and WT littermates. Mice were injected with AAV1/2-Syn1-mRuby2-P2A-GCaMP6s (Rose et al., 2016), which allows functional imaging along with morphological labeling of the neurons (Fig. 1C-D). Because of the rapid disease progression in this mouse line, we performed virus injections and cranial window implantations at the age of 3.5 weeks (Fig. 1A). Two-photon imaging sessions of 15 min each were started 3 weeks after the surgery and carried out at weekly intervals between 6.5 and 9.5 weeks of age (Fig. 1A). Populations of 100-200 L2/3 neurons were imaged in awake head-restrained animals during voluntary locomotion in the dark on a spherical treadmill restricted to movement around one axis (Dombeck et al., 2007) (Fig. 1E). The experiments ended before the animals developed overt neurological symptoms. During the imaging sessions, mice exhibited spontaneous running behavior, which was recorded by an IR-sensitive video camera and/or by an optical mouse sensor (Fig. 1E).

### Increased neuronal activity in HD mice before disease onset

Calcium responses were observed in many L2/3 neurons, with activity remaining quite stable during the imaging period in WT mice, but increasing on average in R6/2 mutants (Fig. 2A-B). Accordingly, the average frequency of calcium transients stayed unchanged between the imaging sessions in WT mice, while it increased in R6/2 animals starting from 8.5 weeks of age (WT, 1612 neurons from 6 mice; R6/2, 2589 neurons from 5 mice; Repeated measures ANOVA, Genotype: F(1, 12597) = 2.01, p = 0.16; Age: F(3, 12597) = 82.91, p < 0.001; Interaction F(3, 12597) = 101.2, p < 0.001; Fig. 2C). The change in transient frequency was also evident when we only analyzed the periods during which the animal remained stationary (only experiments with a minimum of 1% stationary time were included in the analysis; WT, 1346 neurons from 6 mice; R6/2, 2589 neurons from 5 mice; Repeated measures ANOVA, Genotype: F(1, 11599) = 0.08, p = 0.78; Age: F(3, 11599) = 73.43, p < 0.001; Interaction F(3, 11599) = 102.66, p < 0.001; Fig. 2D), indicating that the elevated activity was independent of locomotion. Next, we quantified the fraction of active cells (exhibiting at least one calcium transient during an imaging session, see Materials and Methods) at different time points. This analysis revealed a higher fraction of active cells in R6/2 mice from 8.5 weeks onwards (WT, 7 FOVs from 6 mice; R6/2, 10 FOVs from 5 mice; Repeated measures ANOVA, Genotype: F(1, 45) = 1.11, p = 0.31; Age: F(3, 45) = 3.84, p = 0.02; Interaction F(3, 45) = 3.73, p = 0.02; Fig. 2E). After the last awake imaging session, mice were also imaged under isoflurane anesthesia. In agreement with previous studies, the fraction of active cells was lower in anesthetized compared to awake animals (Greenberg et al., 2008). However, consistent with our findings in awake animals, we observed more active cells in R6/2 mice than in WT littermates (WT, 1612 neurons from 6 mice; R6/2, 2589 neurons from 5 mice. Wilcoxon rank-sum test, p=0.038; Fig. 2F). In summary, these data demonstrate elevated neuronal activity in the cortex of HD mice before the onset of neurological and motor impairments.

**Figure 2.**
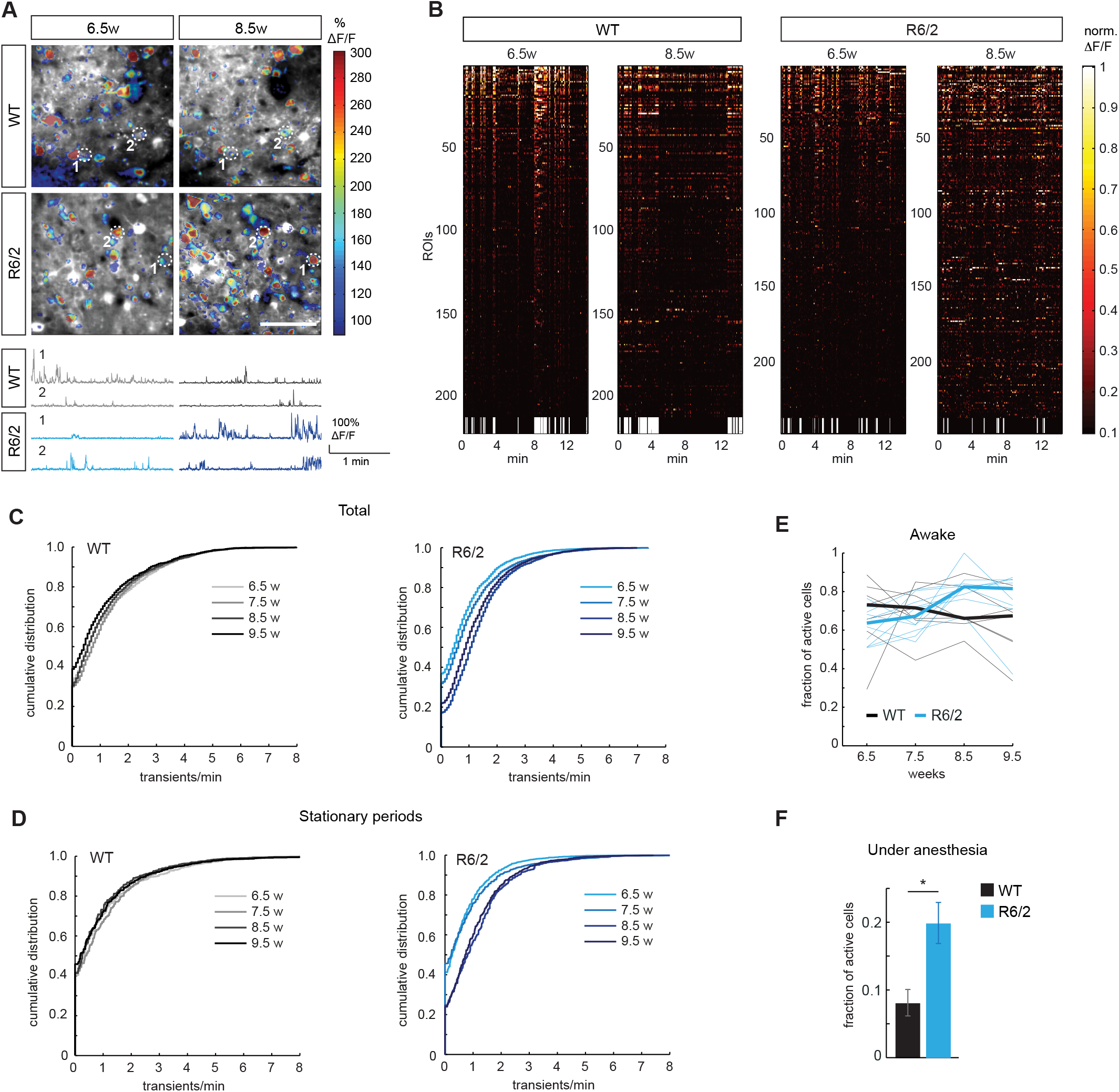
Increased neuronal activity before disease onset in R6/2 mice. **A**, Top: Examples of imaged areas in WT and R6/2 cortex superimposed by activity maps. Color coding shows normalized maximal activity. Bottom: Activity traces of neurons labeled on the images above. **B**, Raster plots of neuronal activity in WT and R6/2 mice during the imaging sessions at 6.5 and 8.5 weeks of age. Cells are sorted by activity at 6.5 weeks. Running episodes are depicted in white and stationary periods in black at the bottom of the plots. **C**, Cumulative distributions of calcium transient frequencies at the indicated time points in WT (left) and R6/2 (right) animals. Note a shift in the distribution towards higher frequencies occurring at 8.5 weeks in R6/2 mice. **D**, Cumulative distributions of calcium transient frequencies during stationary periods at the indicated time points in WT (left) and R6/2 (right) animals. **E**, Fraction of active cells at different imaging time points in WT and R6/2 mice. **F**, Fraction of active cells under isoflurane anesthesia at 9.5 weeks. *p<0.05. Scale bar in a, 50 μm.

### Altered dynamics of single-cell activity in HD mice

We next asked whether individual neurons changed their activity levels during the imaging period and whether the presence of mHTT had an impact on this dynamics. For each of the imaging time points, we quantified the reoccurrence rate of active cells, i.e. the fraction of cells that were active at the given imaging time point and remained active in each of the following imaging sessions. In WT mice, the reoccurrence rates of active cells steadily declined throughout the imaging period, consistent with previous studies describing a high variability of neuronal activity patterns in the motor cortex (Rokni et al., 2007; Peters et al., 2014; Clopath et al., 2017). In R6/2 animals this decline also occurred, but slowed down from 8.5 weeks onwards, resulting in a higher reoccurrence rate compared to WT controls (WT, 7 FOVs from 6 mice; R6/2, 10 FOVs from 5 mice; Repeated measures ANOVA, 6.5 to 9.5 weeks: Genotype: F(1, 45) = 2.61, p = 0.12; Age: F(3, 45) = 130.0, p < 0.001; Interaction F(3, 45) = 3.97, p = 0.01. 7.5 to 9.5 weeks: Genotype: F(1, 30) = 8.59, p = 0.01; Age: F(2, 30) = 94.42, p < 0.001; Interaction F(2, 30) = 5.33, p = 0.01. 8.5 to 9.5 weeks: Genotype: F(1, 15) = 2.01, p = 0.18; Age: F(1, 15) = 94.42, p < 0.001; Interaction F(1, 15) = 2.01, p = 0.18; Fig. 3A). These results suggest that the increase in neuronal activity detected in R6/2 mice is at least partially due to more cells staying active between the imaging sessions.

**Figure 3.**
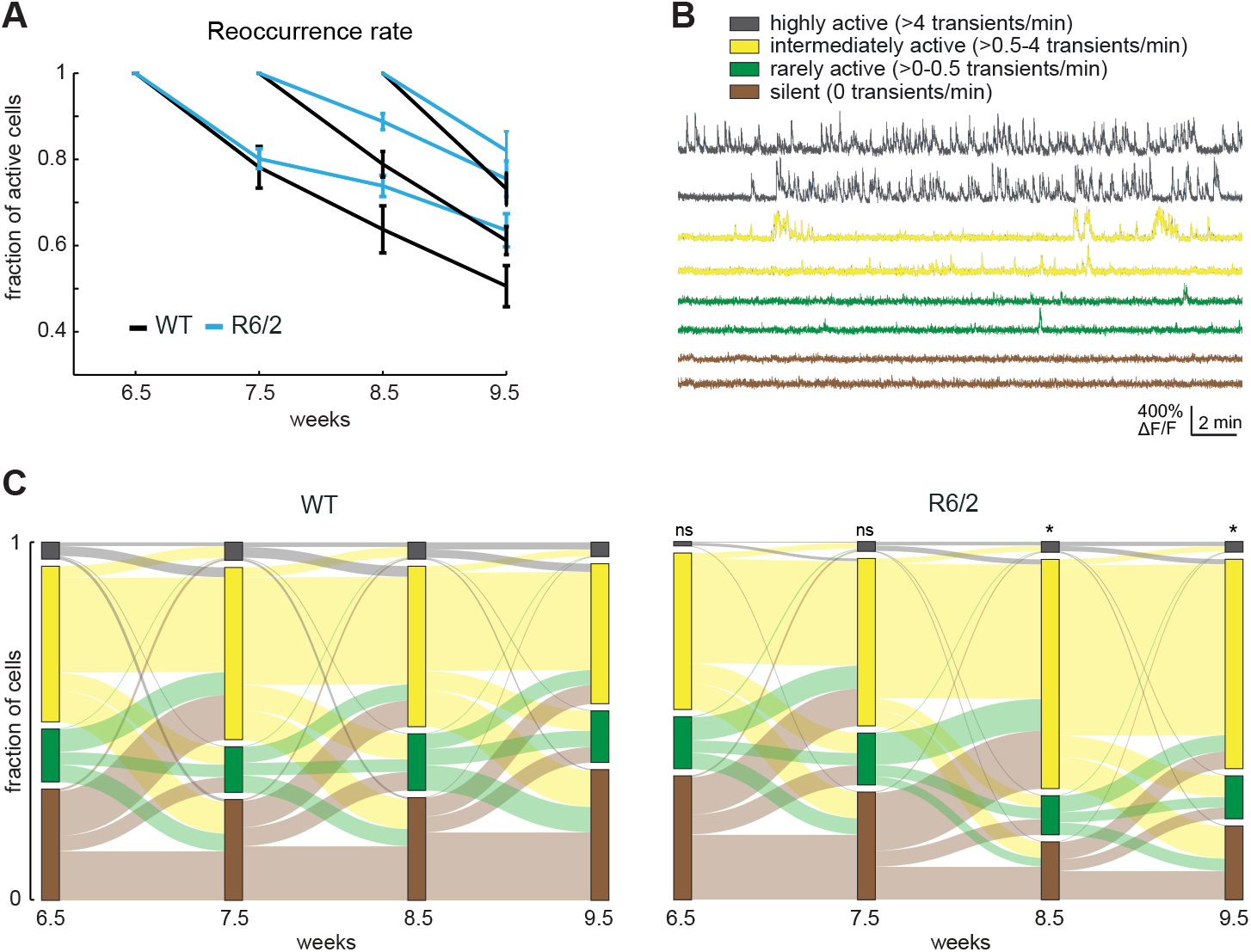
Altered dynamics of single-cell activity. **A**, Reoccurrence rate of active cells in WT and R6/2 mice. **B**, Example traces of highly active, intermediately active, rarely active and silent neurons. **C**, Alluvial plots showing changes between the four activity categories at different imaging time points in WT (left) and R6/2 (right) animals. *p<0.05.

We further examined activity levels of single neurons by subdividing the imaged cells into four activity categories: silent (0 Ca^2^+ transients/min), rarely active (>0-0.5 transients/min), intermediately active (>0.5-4 transients/min), and highly active (>4 transients/min) (Fig. 3B). In the first two imaging sessions, the distribution of cells into these four categories was not significantly different between R6/2 and WT mice (Fig. 3C). However, a clear shift in the distribution was observed in the R6/2 animals between 7.5 and 8.5 weeks of age (WT, 1612 neurons from 6 mice; R6/2, 2589 neurons from 5 mice; Pearson’s Chi-square test, R6/2 vs. WT at 6.5 weeks, p = 0.5347; at 7.5 weeks, p = 0.901; at 8.5 weeks, p = 0.0495; at 9.5 weeks, p=0.0198), which was due to an increase in the fraction of intermediately active cells, as well as reduction in the fraction of silent cells, while the rarely active and the highly active fractions did not change significantly (Fig. 3C). The increase in the intermediately active category in the R6/2 mice at 8.5 weeks was to a large extent due to cells that were classified as silent at 7.5 weeks (26% of silent cells became intermediately active in WT vs. 54% in R6/2; Exact binomial test, p = 1.798e-07). Taken together, these results indicate that activity dynamics are altered in R6/2 mice. The observed increase in activity can at least in part be attributed to more silent cells changing into the intermediately active category, as well as more cells staying active between the imaging sessions.

### Downregulation of synaptic proteins in R6/2 cortex before disease onset

To gain insight into the potential molecular mechanisms underlying the dysregulated activity in the cortex, we took advantage of the spatiotemporally resolved mass spectrometry-based proteomic dataset from R6/2 mice and WT littermates that we obtained previously, including changes in the soluble proteome and composition of insoluble mHTT inclusion bodies (Hosp et al., 2017). For the present study, we focused on cortical samples from 5-week-old (early presymptomatic) and 8-week-old animals (just before the onset of neuronal activity changes). Principal component analysis (PCA) revealed a clear separation of soluble cortical samples of 8-week-old R6/2 mice from the samples of 5-week-old R6/2 and all WT animals (Fig. 4A). We next asked which proteins accounted for this separation. Interestingly, the largest functional group among the main PCA drivers that were downregulated in 8-week-old R6/2 mice were synapse-related proteins (28%, 7 out of 25 proteins; the fraction of synapse-related proteins in the total proteome was 7%, 703 out of 9937 proteins) (Fig. 4B-C, Extended Data Table 4-1). In contrast, no synaptic proteins were found among the main PCA drivers upregulated in R6/2 mice (0 out of 25; Fig. 4B).

**Figure 4.**
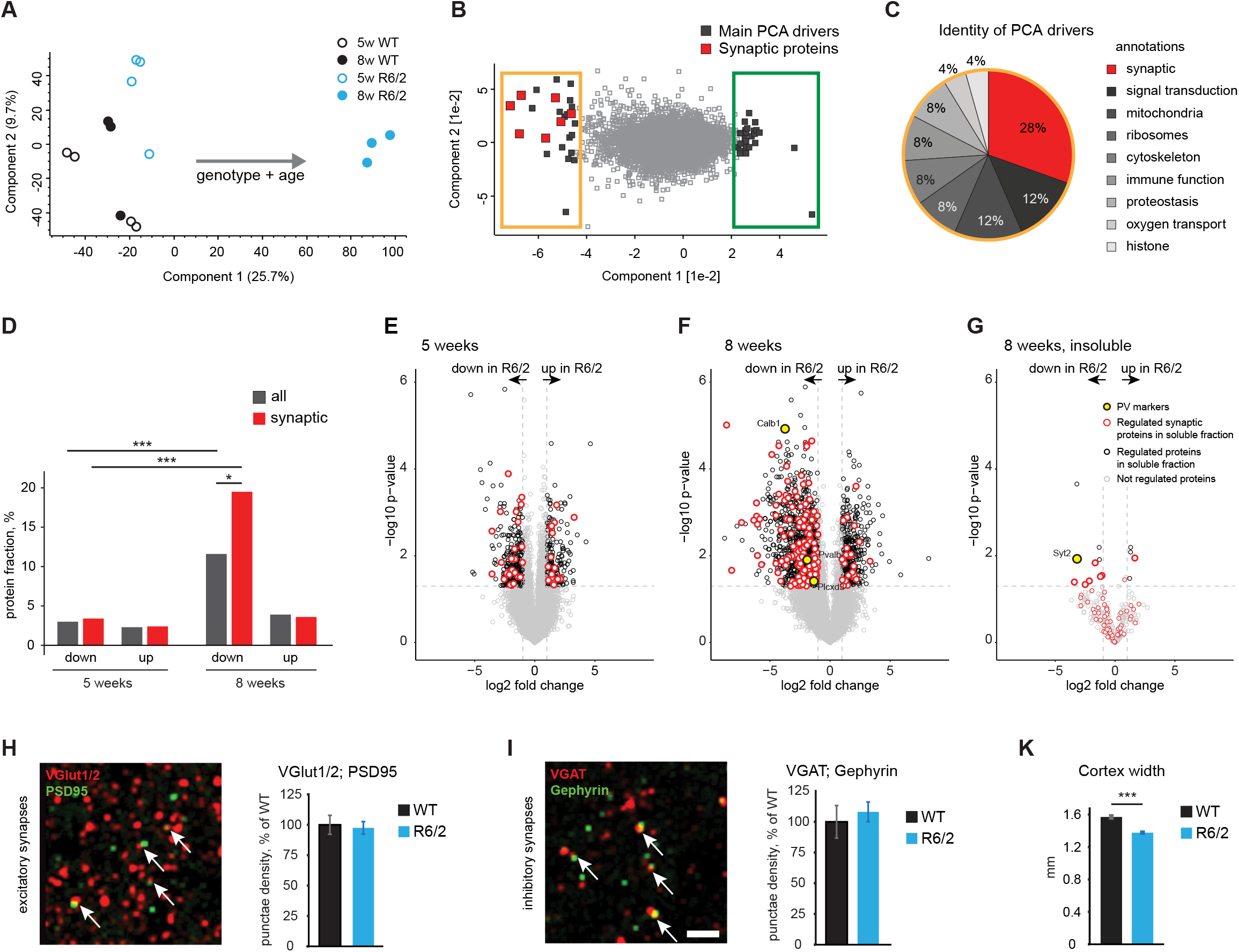
Downregulation of synaptic proteins in R6/2 cortex. **A**, PCA projections of soluble cortical samples from 5- and 8-week old R6/2 mice and WT littermates. **B**, Main PCA drivers of the separation that are downregulated (yellow frame) and upregulated (green frame) in 8-week-old R6/2 mice. Synapse-related proteins are highlighted in red. Main drivers were defined as the top 25 proteins accounting for the separation of samples in the PCA. **C**, Functional groups of the main PCA drivers downregulated in 8-week-old R6/2 mice (indicated by the yellow frame in B). See also Table 4-1. **D**, Fraction of proteins significantly up- or downregulated in the soluble fraction of the R6/2 cortex at the indicated ages. See also Table 4-2. **E-F**, Volcano plots showing proteins significantly up- or downregulated in the soluble fraction of 5-week-old (E) and 8-week-old (F) R6/2 cortex. Empty black circles, all significantly regulated proteins; empty red circles, significantly regulated synaptic proteins. **G**, Volcano plot of the cortical insoluble fraction at 8 weeks, showing only proteins significantly downregulated in the soluble proteome at this age (see F). See also Table 4-3. Significantly downregulated PV cell markers are shown as filled yellow circles and labelled on the plots in F and G. **H-I**, Representative images (left) and quantification (right) of excitatory (H) and inhibitory (I) synapses in the L2/3 of the M1 area at 8 weeks. Synapses were identified by the overlap or close apposition (arrows) of a presynaptic (red) and postsynaptic (green) marker. **K**, M1 cortex width in 8-week-old R6/2 and WT mice. *p<0.05; ***p<0.001. Scale bar in I, 3 μm.

While there were no major changes in the soluble cortical proteome of 5-week-old mice, a global downregulation of multiple proteins in the soluble fraction occurred at 8 weeks of age in the R6/2 cortex (Fig. 4D-F), consistent with the data from other brain regions (Hosp et al., 2017). At 8 weeks of age, we also observed a pronounced decrease in synaptic protein levels (Fig. 4F). Importantly, the fraction of downregulated synaptic proteins (19%, 137 out of 703 proteins) was significantly higher than the downregulated fraction of the total proteome (12%, 1153 out of 9938 proteins) (Exact binomial test, p=0.0434; Fig. 4D and Extended Data Table 4-2), pointing towards a specific loss of synaptic proteins that is not merely due to the general remodeling of the whole proteome. We next asked if this downregulation of synaptic proteins might be due to their sequestration within mHTT inclusions. Out of 137 synaptic proteins reduced in the soluble fraction at 8 weeks, 60 were detected in the insoluble proteome, which largely consists of mHTT inclusions. Remarkably, only 1 of the detected proteins was significantly increased in R6/2 mice, while 6 were decreased, and 53 not significantly altered (Fig. 4G, Extended Data Table 4-3). Sequestration within mHTT inclusions is therefore likely not a major mechanism of synaptic protein decrease at an early stage of HD progression. This is distinct from the situation in 12-week-old R6/2 brains, where many proteins downregulated in the soluble pool are enriched in the insoluble material, indicative of their recruitment to mHTT inclusion bodies (Hosp et al., 2017). In summary, our proteomic results reveal a broad downregulation of synaptic components already before disease onset.

### Loss of PV-positive inhibitory inputs onto PCs

Lower levels of synaptic proteins in the R6/2 mice might reflect a loss of synapses. A decrease in synapses in the R6/2 cortex at an advanced disease stage has been reported previously (Murmu et al., 2015). However, it has remained unclear whether synapse numbers are already reduced before disease onset. We therefore performed immunostainings for excitatory and inhibitory pre- and postsynaptic markers in L2/3 of the M1 cortex of 8-week-old animals and quantified densities of opposing VGlut1/2; PSD-95 puncta as putative excitatory synapses and opposing VGAT; Gephyrin puncta as putative inhibitory synapses (Fig. 4H-I). These experiments did not reveal significant changes in total excitatory or inhibitory synapse densities at 8 weeks (3 FOVs per mouse from 5 WT and 4 R6/2 mice. Student’s t-test; excitatory synapses, p=0.8279; inhibitory synapses, p=0.6773; Fig. 4H-I). However, since cortical thickness is already significantly reduced in R6/2 mice at this age (WT, 5 mice, R6/2, 4 mice; Student’s t-test, p=0.0003; Fig. 4K), the overall number of synapses is likely to be decreased. While higher-resolution methods will be required for a more precise quantification, our current results therefore suggest that both excitatory and inhibitory synapse numbers are reduced in R6/2 mice already before disease onset.

We next focused on a more specific subset of synapses, PV-positive IN terminals on PCs. PV cells represent the most numerous IN subtype in the cortex, form synaptic terminals on the perisomatic region of their target cells and constitute the strongest source of inhibition onto PCs (Pfeffer et al., 2013; Tremblay et al., 2016). Interestingly, we found several specific markers of PV INs, including Pvalb, Calb1, and Plcxd3 (Tasic et al., 2016; Tasic et al., 2017; Mayer et al., 2018), to be downregulated in the soluble cortical proteome of R6/2 mice, as well as Synaptotagmin 2, a specific marker of PV synaptic terminals (Sommeijer and Levelt, 2012), to be downregulated in the insoluble fraction (Fig. 4F-G). Immunostainings revealed a significant reduction in the area of PV+ terminals surrounding NeuN-labeled PC cell bodies in L2/3 of the M1 cortex of R6/2 mice at 8 weeks, but not at 5 weeks of age (WT, 74 PCs from 5 mice and 63 PCs from 5 mice for 5 and 8 weeks of age, respectively; R6/2, 74 PCs from 5 mice and 58 PCs from 4 mice for 5 and 8 weeks of age, respectively; Student’s t-test, 5 weeks, p=0.3131; 8 weeks, p=0.0257. Fig. 5A-B).

**Figure 5.**
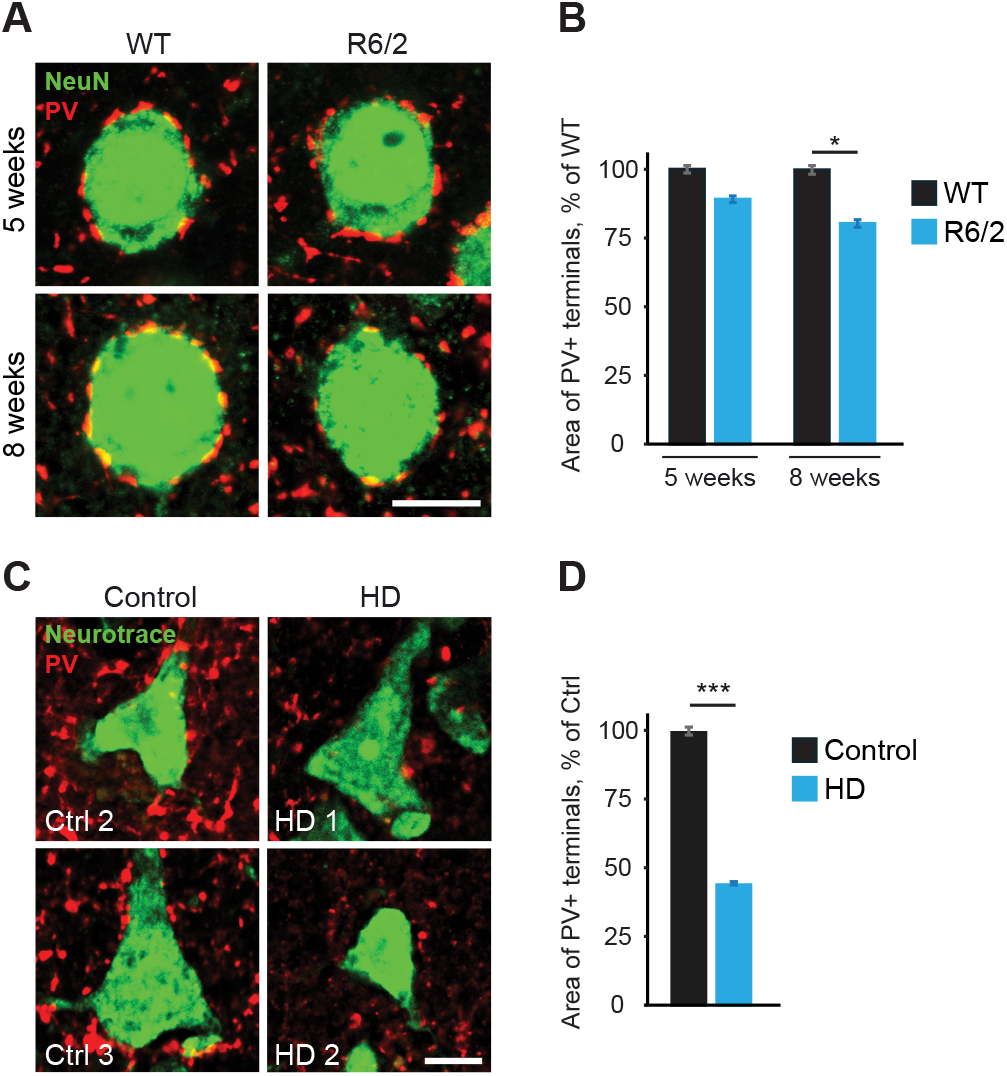
Reduction in inhibitory inputs from PV INs in R6/2 and HD patient cortex. **A-B**, Representative images (A) and quantification (B) of PV synaptic terminals (red) on NeuN-labeled PC cell bodies (green) in L2/3 of M1 cortex from R6/2 and WT controls. **C-D**, Representative images (C) and quantification (D) of PV synaptic terminals (red) on Neurotrace-labeled PC cell bodies (green) in L2/3 of M1 cortex from HD autopsy cases and controls. Scale bars in A and **C**, 10 μm.

To validate these histological findings in human disease cases, we performed immunostainings for PV+ terminals in postmortem brain tissue of HD patients and age-matched controls. Consistent with the results from R6/2 mice, we observed a marked reduction in the area of PV+ puncta around L2/3 PCs in the primary motor cortex (Ctrl, 69 PCs from 3 brains; HD, 75 PCs from 3 brains; Student’s t-test, p=1.436e-15. Fig. 5C-D). In summary, loss of PV+ inhibitory inputs onto PCs occurs both in R6/2 mice and in human HD cases, suggesting that weakened inhibition from PV+ INs might at least partially explain dysregulated cortical activity in HD.

## Discussion

Using chronic *in vivo* imaging in awake animals, we demonstrate an overall increase in cortical activity in HD mice. While our results are in agreement with previous reports describing dysregulation of activity and higher excitability in rodent models of HD (Walker et al., 2008; Cummings et al., 2009; Miller et al., 2011), we were able for the first time to observe these changes at a single-cell resolution, in awake animals already at a presymptomatic stage. Our findings therefore suggest that impaired cortical activity might critically contribute to disease onset. We also show that the excessive activity is not related to voluntary locomotion. This activity might therefore be aberrant, leading to higher noise in the cortical networks and thereby degrading information processing, as observed previously in a mouse model of Alzheimer’s disease (Liebscher et al., 2016).

Our chronic imaging approach further revealed altered activity dynamics at single-cell level, with more neurons remaining active between imaging sessions, as well as more cells changing from silent to intermediately active. Highly dynamic activity patterns that are characteristic of the motor cortex and believed to be important for motor learning (Peters et al., 2014) therefore appear to be compromised in these mice. These findings provide a possible mechanism for the motor learning impairments observed in HD mouse models (Trueman et al., 2007; Woodard et al., 2017) as well as HD patients (Heindel et al., 1988).

Dysfunction of cortical circuitry might in turn lead to profound changes of neuronal function in the striatum. Early in HD progression, hyperactivity of striatal neurons was described R6/2 mice (Rebec et al., 2006), and elevated glutamate levels were detected in the striatum of HD patients (Taylor-Robinson et al., 1996). These alterations could in part be due to increased activation of the striatum by cortical inputs. Remarkably, removal of cortical afferents resulted in amelioration of HD phenotypes in R6/2 mice (Stack et al., 2007), and expression of mHTT in cortical neurons was necessary and sufficient to cause functional impairments of the striatal compartment in a corticostriaral network reconstructed *in vitro* (Virlogeux et al., 2018). It remains to be tested whether restoration of normal cortical activity levels, e.g. with the help of chemogenetic tools, would also be sufficient to rescue or delay HD symptoms.

Searching for the underlying circuit mechanisms of impaired cortical activity, we observed a reduction in inhibitory PV+ inputs onto PCs in R6/2 mice, which we confirmed in HD autopsy cases. These findings suggest that abnormalities of cortical function in HD could at least in part be explained by weakened inhibition. Reduced frequency of inhibitory postsynaptic currents has been described in L2/3 PCs in the motor and somatosensory cortex in several HD mouse models at a symptomatic stage (Gu et al., 2005; Spampanato et al., 2008; Cummings et al., 2009). Consistently, human studies with the use of transcranial magnetic stimulation also revealed abnormal cortical excitability due to dysfunctional inhibition in presymptomatic and early-stage HD patients (Schippling et al., 2009; Philpott et al., 2016), emphasizing the translational value of our findings.

Several previous reports suggested that PV cells might play a role in HD pathogenesis. PV INs develop mHTT inclusion bodies and show electrophysiological changes in HD mouse models (Meade et al., 2002; Spampanato et al., 2008). Selective expression of mHTT in this IN population results in specific HD-related phenotypes, including impaired cortical inhibition (Dougherty et al., 2014). It should, however, be noted that our results do not exclude a possible involvement of other IN subtypes in the cortical HD pathology. With the help of the available genetic tools for labeling specific IN subpopulations and manipulating their activity *in vivo* (Taniguchi et al., 2011), in future studies it should be possible to uncover the precise roles of particular IN subtypes in this disease. Interestingly, impairments in the function of inhibitory neurons have also been suggested to underlie neural circuit defects in mouse models of Alzheimer’s disease (Verret et al., 2012; Busche and Konnerth, 2016). Excitation/inhibition imbalance and resulting dysregulation of cortical activity may therefore represent common phenomena in various neurodegenerative disorders.

What might be the molecular link between expression of mHTT and defects of neuronal communication in the cortex? Our proteomic analyses demonstrated a major downregulation of synaptic proteins in R6/2 mice. In addition to confirming reductions in synaptic proteins shown at advanced disease stages in HD mice and human brain samples (Morton and Edwardson, 2001; Morton et al., 2001; Smith et al., 2007; Skotte et al., 2018), our dataset reveals a broad downregulation of synaptic components that occurs already before disease onset. Surprisingly, the majority of synaptic proteins that were downregulated in the soluble fraction were not altered in the insoluble material at 8 weeks of age, arguing against sequestration as a major mechanism of synaptic protein depletion at this disease stage (Murmu et al., 2013; Kim et al., 2016; Hosp et al., 2017). One likely mechanism of synaptic protein reduction might be transcriptional dysregulation, as genes related to synaptic signaling were among the most prominent dysregulated gene clusters in transcriptomic analyses of HD mice and human iPSC-derived neural cultures (Tang et al., 2011; Langfelder et al., 2016; HD iPSC Consortium, 2017). While synapse loss and downregulation of synaptic proteins might be expected to go along with a reduction in neuronal activity, we and others (Cummings et al., 2009) in fact observed the opposite effect in the cortex of HD mice. This underscores the complexity of HD pathogenic mechanisms and the value of *in vivo* imaging studies in deciphering cortical microcircuit alterations in disease models. Taken together, our chronic single-cell resolution imaging approach, in combination with state-of-the-art proteomic analyses and histological experiments, points towards disturbed cortical excitation/inhibition balance preceding disease onset as a key mechanism in HD pathogenesis.

## Author Contributions

J.N. and E.K.S.-T. acquired the in vivo imaging data. E.K.S.-T. performed the histological experiments. S.G.-A. conducted the behavioral tests. J.N., E.K.S.-T. and S.L. performed data analysis. F.H. and M.M. provided unpublished proteomic data. T.A. provided the human tissue and gave advice on neuropathological analyses. R.K., S.L. and I.D. supervised the project. J.N., R.K., S.L. and I.D. wrote the paper.

## Acknowledgements

We thank Nejc Dolensek for the implementation of the speed sensor; Narasimha Reddy Vaka, Robert Kasper, Diego Sangineto and Henry Haeberle for technical support with microscope hardware; Pieter Goltstein and Tobias Rose for providing custom-made tools for awake imaging; Hakan Kucukdereli and Pieter Goltstein for insightful discussions; Stefanie Huschenbett, Mario Rivera-Cortes and Eneyda Kowalski for excellent animal care; Julia Boshart for excellent assistance with histology; André Wilke for quantification of immunostainings; André Wilke and Tammo von Knoblauch for mouse genotyping; and Dominique Förster for critically reading the manuscript. This work was funded by the European Research Council Synergy Grant under FP7 GA number ERC-2012-SyG_3í8987-Toxic Protein Aggregation in Neurodegeneration (ToPAG) to R.K. and M.M., and by the Max Planck Society for the Advancement of Science.

## Extended data

**Extended Data Table 4-1**. Main PCA drivers for the soluble cortical proteome.

**Extended Data Table 4-2**. Soluble cortical proteome of R6/2 mice at 5 and 8 weeks.

**Extended Data Table 4-3**. Insoluble fraction of R6/2 cortical samples at 5 and 8 weeks.

## Notes

**Conflicts of Interest** The authors declare no conflicts of interest.

